# Retrotransposons are specified as DNA replication origins in the gene-poor regions of Arabidopsis heterochromatin

**DOI:** 10.1101/090183

**Authors:** Zaida Vergara, Joana Sequeira-Mendes, Jordi Morata, Elizabeth Hénaff, Ramón Peiró, Celina Costas, Josep M. Casacuberta, Crisanto Gutierrez

## Abstract

Genomic stability depends on faithful genome replication. This is achieved by the concerted activity of thousands of DNA replication origins (ORIs) scattered throughout the genome. In spite of multiple efforts, the DNA and chromatin features that determine ORI specification are not presently known. We have generated a high-resolution genome-wide map of ORIs in cultured Arabidopsis thaliana cells that rendered a collection of 3230 ORIs. In this study we focused on defining the features associated with ORIs in heterochromatin. We found that while ORIs tend to colocalize with genes in euchromatic gene-rich regions, they frequently colocalize with transposable elements (TEs) in pericentromeric gene-poor domains. Interestingly, ORIs in TEs associate almost exclusively with retrotransposons, in particular, of the Gypsy family. ORI activity in retrotransposons occurs independently of TE expression and while maintaining high levels of H3K9me2 and H3K27me1, typical marks of repressed heterochromatin. ORI-TEs largely colocalize with chromatin signatures defining GC-rich heterochromatin. Importantly, TEs with active ORIs contain a local GC content higher than the TEs lacking them. Our results lead us to conclude that ORI colocalization with TEs is largely limited to retrotransposons, which are defined by their transposition mechanisms based on transcription, and they occur in a specific chromatin landscape. Our detailed analysis of ORIs responsible for heterochromatin replication has also implications on the mechanisms of ORI specification in other multicellular organisms in which retrotransposons are major components of heterochromatin as well as of the entire genome.

## Introduction

Reliable and complete genome duplication is crucial to maintain genomic stability. In eukaryotes, DNA replication occurs during the S-phase of the cell cycle and is initiated at multiple genomic locations, known as DNA replication origins (ORIs). Over the past years, detailed genome-wide maps of ORIs have been generated for various multicellular organisms such as cultured Drosophila, mammalian and Arabidopsis cells (Sanchez et al. 2012; Mechali et al. 2013; Renard-Guillet et al. 2014; Comoglio et al. 2015). ORI specification and activation depends on several variables, including the cell’s type and the physiological state as well as specific chromatin features, frequently including those associated with open chromatin (MacAlpine and Almouzni 2013; Mechali et al. 2013; Sequeira-Mendes and Gutierrez 2015). A preference of ORIs for colocalizing with genic regions, in particular highly expressed genes, seems to be a common observation across all organisms studied so far (Costas et al. 2011; Lubelsky et al. 2014; Cayrou et al. 2015; Sequeira-Mendes and Gutierrez 2015).

Chromatin can be divided into heterochromatin, which is densely compacted for most of the cell cycle, and euchromatin, with a relatively less dense organization. Genes are not evenly located throughout the chromosomes, as they are more frequent in the euchromatic chromosome arms. This distribution is the inverse of that of transposable elements (TEs), which tend to accumulate in heterochromatic domains (Bennetzen and Wang 2014). In Arabidopsis, several TE families account for 21% of the genome and, although some of them are scattered along chromosome arms, most TEs concentrate in the pericentromeric heterochromatin (Ahmed et al. 2011; Feng and Michaels 2015). Whilst previous studies have reported the link between DNA replication fork progression and the establishment of heterochromatin (Nikolov and Taddei 2016), the genomic features that contribute to specify ORIs in heterochromatin have not been studied and, consequently, are very poorly understood.

Here we have used Arabidopsis cultured cells to study in detail the genomic features defining ORI localization in heterochromatin, largely concentrated in the pericentromeric regions. We found that whereas in euchromatic chromosome arms the vast majority of ORIs (94.9%) colocalize with genes, in the pericentromeric gene-poor regions TEs contribute a significant fraction of ORIs (33.7%). Our study also shows that not all TEs serve equally as ORIs. Retrotransposons, and in particular Gypsy elements, more frequently colocalize with them. Furthermore, we found that a specific chromatin landscape mainly characterized by a GC-rich heterochromatic state, is a determinant feature for ORI localization in heterochromatin. Together, our findings suggest that the characteristics of the chromatin associated to each family of TEs, their genomic organization and the retrotransposons’ potential for transcription are key to determine their capacity to contain ORIs. Our study serves the basis to tackle in the future the question of how the ORI specification and replication machineries gain access to the highly compact heterochromatic regions to achieve its duplication during S-phase.

## Results

### High-resolution identification of ORIs in transposable elements

One of the strategies to identify ORIs relies on the isolation of small newly synthesized DNA molecules from replication bubbles. The identification of ORIs responsible for replication of pericentromeric heterochromatin requires very reliable genome annotation and peak calling algorithms. Probably because of that, it has never been undertaken systematically. In the case of *Arabidopsis thaliana,* an updated genome annotation (TAIR10), including highly repetitive pericentromeric regions, is now available. Also, various peak calling algorithms have been reported, among which we found that MACS1.4 (Zhang et al. 2008) is well suited for our purpose as it detects peaks of relatively small size (<0.5-1kb) and of low representation in the sample.

We have used these tools and sequencing data of purified BrdU-pulsed DNA extracted from Arabidopsis cultured cells (GSE21828) to generate a high-resolution map of ORIs, paying particular attention to those located in heterochromatic regions. Genes and TEs in Arabidopsis are not homogenously distributed along the chromosomes. TEs are largely, although not exclusively, concentrated in heterochromatic domains, and in particular at pericentromeric regions, whereas most genes are located in non-pericentromeric euchromatin domains (Ahmed et al. 2011). Since heterochromatin domains contain highly repetitive sequences, such as TEs, we first concentrated in hits that unequivocally aligned to only one genomic location, leaving multihit reads for a subsequent analysis. This approach obviously rendered an underestimation of ORIs mapping to these regions but it provided a more confident dataset of ORIs responsible for heterochromatin replication (Fig. 1A).

**Figure 1.**
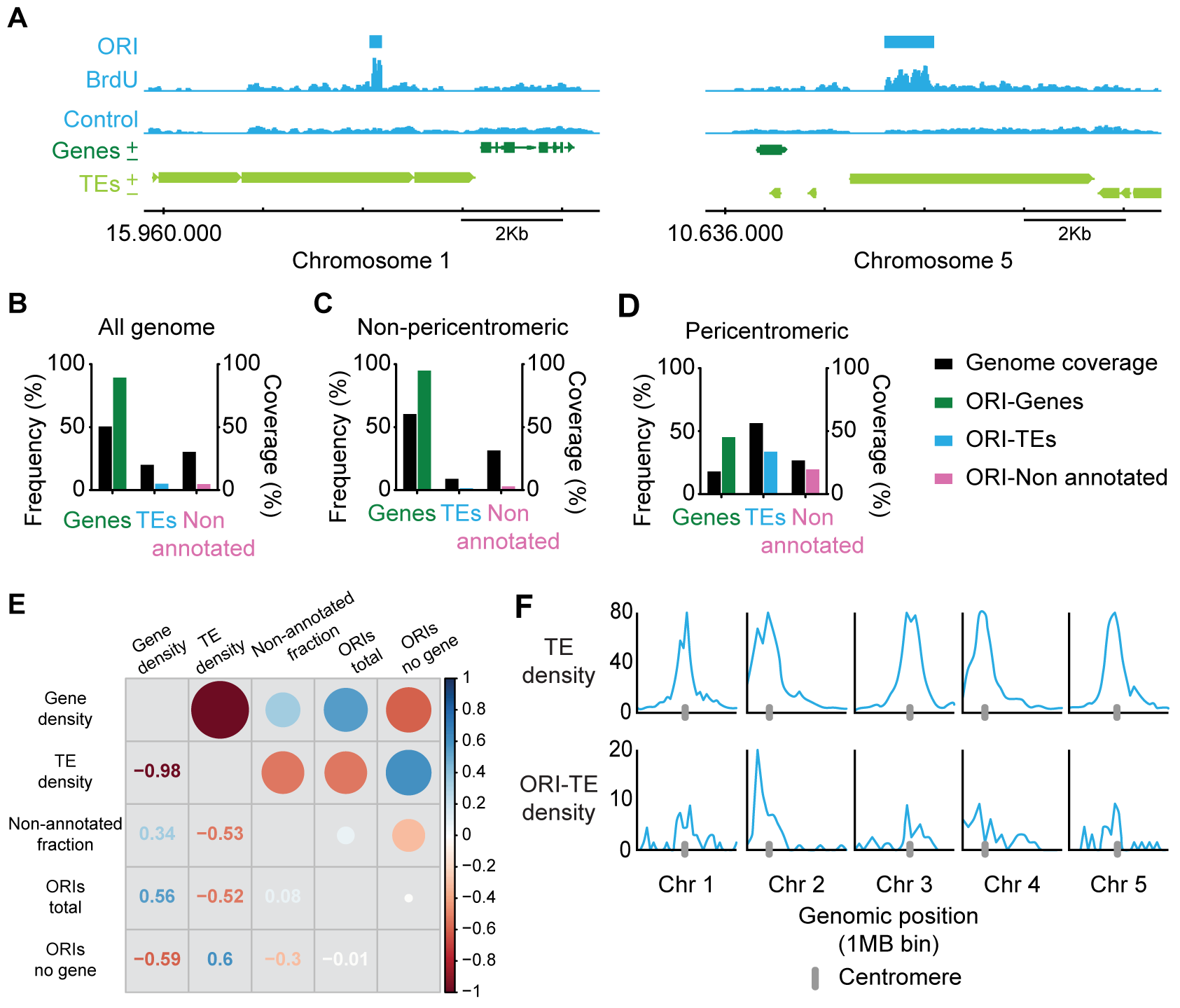
Genomic location of Arabidopsis DNA replication origins. *(A)* Representative genome-browser views of regions containing ORIs of chromosomes 1 and 5, as indicated. BrdU-peaks defining ORIs relative to the control are indicated (light blue bars). Genes (dark green) transcribed from each strand and TEs (light green) are shown along the chromosome together with the coordinate scale. Fraction of ORIs found in genes, TEs and non-annotated regions in (*B*) all the Arabidopsis genome, (*C*) the non-pericentromeric regions and (*D*) the pericentromeric regions, defined as having a gene frequency ≤40%, shown with the respective genome coverage. (*E*) Overall correlation between gene, TE and non-annotated fraction coverage and total ORIs and ORIs not located in genes. Correlations are represented with circles (gradation of red, anticorrelation; gradation of blue, positive correlation). The size of the circles corresponds to the correlation coeficient, also indicated in the other half of the plot. (*F*) TE density (% of nucleotides in TEs per 1 Mb bin) (upper panels) and chromosomal distributions of ORI-TEs across the five Arabidopsis chromosomes (lower panels).

Our analysis showed that ORIs have a strong preference to colocalize with genes. Out of a total of 3230 ORIs in the entire genome (Table S1), 2888 (89.4%) colocalized with genes and 161 (4.9%) with TEs (Figure 1B; Fig. S1; Table S2), a result in accordance with our previous overall analyses (Costas et al. 2011). However, this analysis also showed that the proportions change drastically when we consider separately non-pericentromeric (chromosome arms) and pericentromeric regions. Indeed, whereas almost all ORIs (94.9%) colocalize with genes in gene-rich domains of chromosome arms, less than half of ORIs (46.7%) colocalize with genes in the pericentromeric gene-poor regions (Fig. 1C,D; Table S2). Furthermore, the distribution of ORIs not located in genes positively correlates with the distribution of TEs, and not with the distribution of non-annotated regions (Fig 1E). Analysis of ORI-TE density along the Arabidopsis chromosomes visualizes the preference of non-genic ORIs to colocalize with TEs in pericentromeric regions (Fig. 1F). These results suggest that TE sequences may be selected as ORIs in regions with a low gene density such as pericentromeric regions. To evaluate if the distribution of ORIs in TEs was affected by choosing the uniquely mapped reads, we repeated the analysis using the multihit sequence reads and found very similar results (Table S3). Also importantly, similar results were obtained using the BayesPeak algorithm (Cairns et al. 2011) (data not shown).

### ORI-TEs preferentially colocalize with retrotransposons

TEs constitute a very heterogeneous type of repetitive elements that can be divided in different classes and families based on their structure and transposition mechanisms (Wicker et al. 2007; Deragon et al. 2008). Therefore, we first asked whether ORIs in TEs were homogenously distributed among the various TE families and found a striking preference for ORIs to associate with certain TE families (Table S4). The vast majority of ORI-TEs (83.9%) is located in retrotransposons of the Gypsy, Copia and LINE families that account only for 42.4% of the TE genome space (Fig. 2A). In particular, Gypsy elements that cover 29.4% of the TE genome space contain ∼50% of all ORI-TEs. On the contrary, ORI-TEs are clearly under-represented in other families, especially of DNA transposons. Helitrons, which have a similar prevalence compared to Gypsys, lack any detectable ORI-TEs and DNA/MuDR that account for 15.7% of the TE genome space contain only 3.4% of all ORI-TEs (Fig. 2A). Since the pericentromeric regions concentrate most ORI-TEs, the tendency of ORI-TEs to colocalize with Gypsy elements could simply be due to the skewed distribution of Gypsy elements towards pericentromeric regions. However, our data show that ORI-TEs are overrepresented in Gypsy elements also in non-pericentromeric regions (Fig. 2B,C).

**Figure 2.**
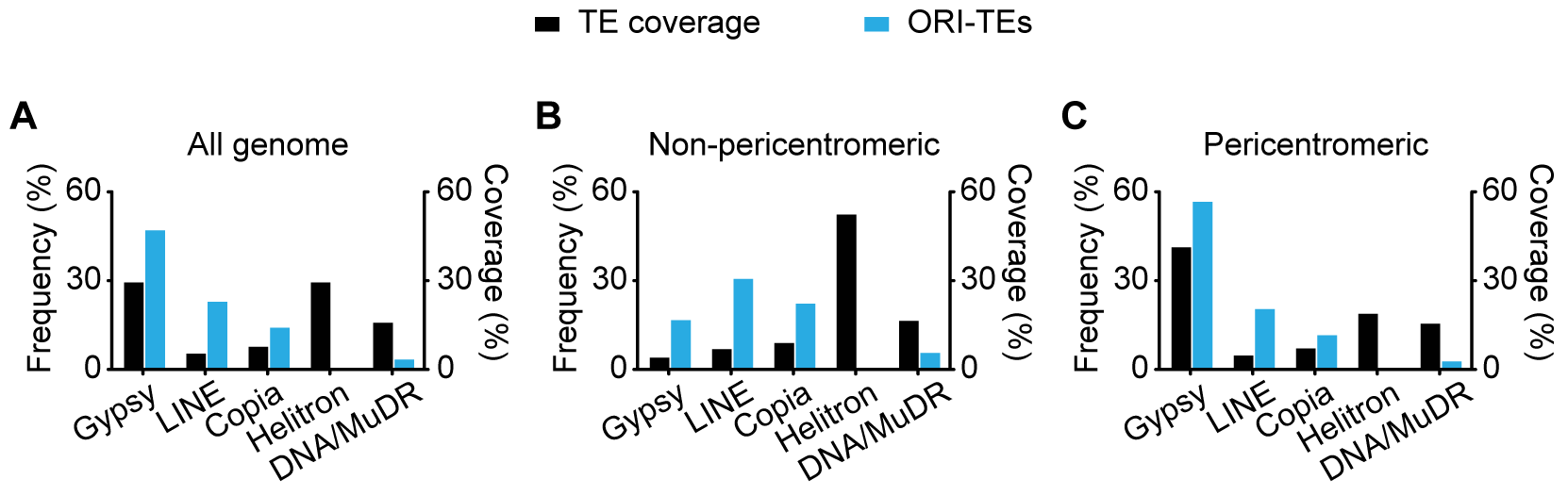
Frequency distribution of ORI-TEs in TE families. *(A)* All the Arabidopsis genome. *(B)* Non-pericentromeric regions. *(C)* Pericentromeric regions, (blue bar) shown with the respective TE family nucleotide coverage of total TE nucleotides (black bar).

Moreover, the complete lack of ORIs in Helitrons, which account for more than 18% of the TEs in pericentromeric regions, also shows that this is not the case. Analysis of the multihit sequences revealed similar results (Fig. S2), indicating that the lack of ORIs in Helitrons is not due to a bias derived from sequence alignments problems. Together, these observations demonstrate that when ORIs associate with TEs they have a significant preference to colocalize with retrotransposons and specifically Gypsy elements, whereas they tend to be excluded from DNA transposons, in particular from Helitron elements.

### Short nascent DNA strands (SNS) enrichment confirms the activity of ORIs mapped by BrdU-seq

To validate our ORI mapping strategy using an independent method we determined the activity of a number of ORIs by quantitative PCR enrichment of a purified sample of short nascent strands (SNS) isolated from DNA replication bubbles (Gerbi and Bielinsky 1997; Cayrou et al. 2012b). For a detailed validation of ORI activity we designed sets of primer pairs across a chromosomal region containing one ORI overlapping with a TE in the arm of chromosome 1 (AT1TE62820) and another ORI ∼70 kb apart, colocalizing with a downstream gene (AT1g51350) within a typical euchromatic region. Cultured Arabidopsis cells were synchronized in G0 by sucrose deprivation and then samples were extracted 2 (G1/S), 3.5 (early S) and 7 h (late S) after release from the sucrose block. qPCR analysis was carried out in two consecutive fractions of the sucrose gradient to ensure reproducibility of the data. As expected, none of the ORIs selected were active at the earliest time point analyzed, 2h after release of the sucrose block (Fig. 3A). At later time points, a clear enrichment was detected in both cases, revealing the activity of these two ORIs in the cell population. Also, it is worth noting that the ORI located within a gene (Fig. 3A, right panels) was ∼5-10-fold more active than the ORI colocalizing with a TE (Fig. 3A, left panels). These experiments confirm that both predicted ORIs, located in a TE and in a gene, indeed function as ORIs. This analysis also showed that an ORI located at a TE in a chromosome arm is active in cultured cells, even when another stronger ORI is in the neighborhood, less than ∼70 kb apart.

**Figure 3.**
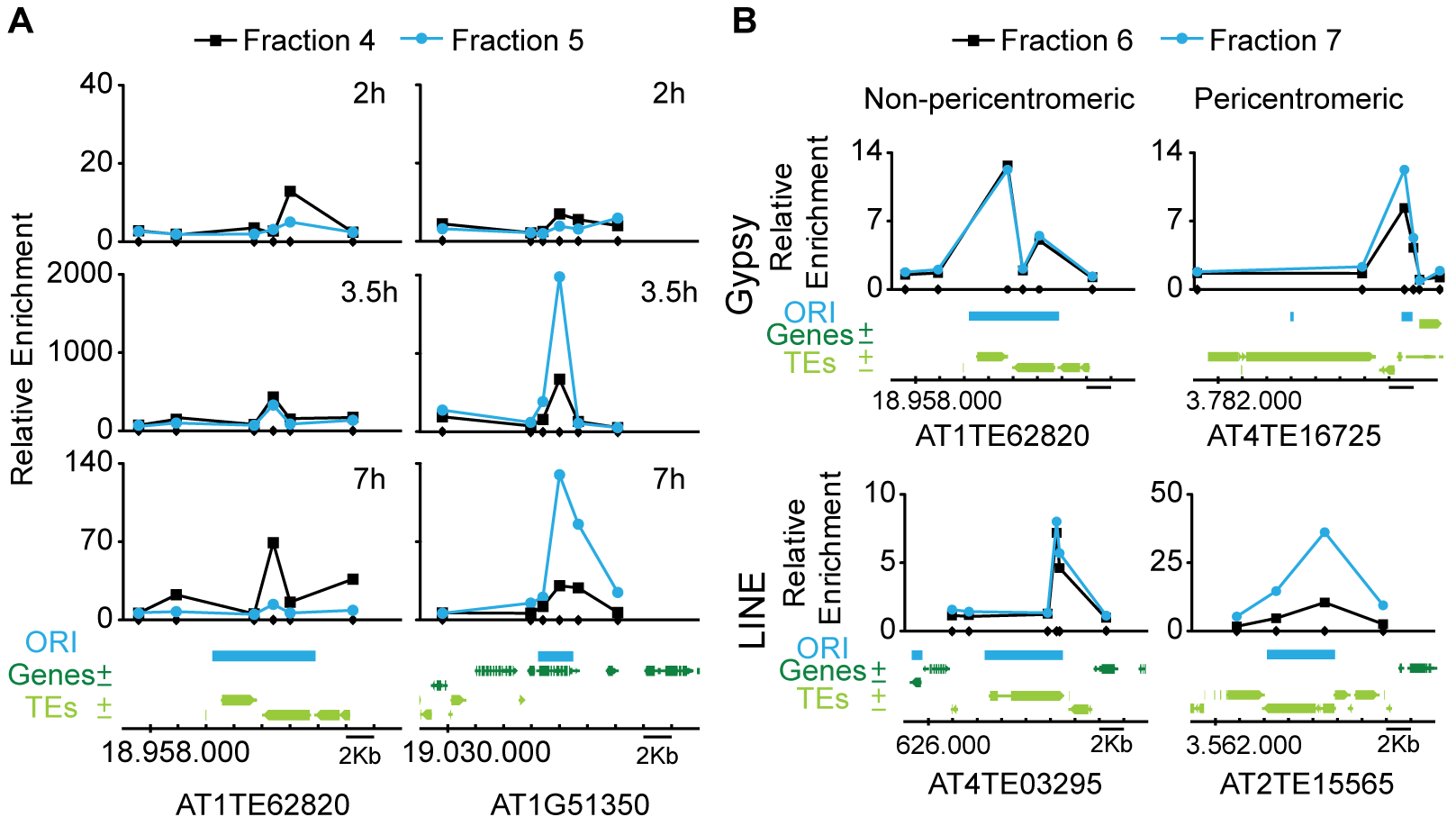
DNA replication origin activity determined by short nascent strand (SNS) abundance by qPCR. *(A)* Measurement of ORI activity in synchronized Arabidopsis MM2d cells at various times after releasing the block, as indicated (2h, G1/S; 3.5h, S; 7h, late S). In each case, the confidence of ORI activity was assessed by analyzing in two biological replicates two consecutive fractions, as indicated at the top. The fractions belong to the same gradient used for purification of SNS and contain DNA molecules ranging 500-2500 bp in size. Two ORI-containing regions (left panels, ORI colocalizing with a TE; right panels, ORI colocalizing with a neighbor gene) were analyzed. The location of primer pairs scanning the region is indicated by small dots on the X-axis. Enrichment values were made relative to the flanking region and normalized against gDNA. The genomic region under study depicting the location of ORI, genes and TEs is at the bottom. Chromosomal coordinates are indicated. *(B)* Measurement of ORI activity in asynchronous Arabidopsis MM2d cell cultures. The ORI-TEs were chosen according to their family (Gypsy and LINE) and location (non-peri- and pericentromeric), as indicated. The location of primer pairs is indicated by small dots on the X-axis. Two consecutive fractions were analyzed, as described for panel *A.* Fractions containing smaller DNA fragments did not give reproducible SNS-qPCR enrichment. Enrichment values were made relative to a negative region that does not content any ORI or TE (AT2G28970). The genomic region under study depicting the location of ORI, genes and TEs is at the bottom. Chromosomal coordinates are indicated

We also wanted to evaluate the activity of different ORI-TEs according to the TE family they colocalize with. Thus, we chose to validate and analyze in asynchronous cells, four genomic regions containing ORI-TEs: two belonging to the Gypsy family and two belonging to the LINE family (where ORIs are highly and moderately over-represented, respectively), and in each case one ORI located in pericentromeric heterochromatin and another in non-pericentromeric heterochromatic patches within the euchromatic arms. These regions were also selected based on the possibility to design a set of primer pairs that unequivocally identify them. We found that all ORI-TEs analyzed here were active as revealed by the qPCR enrichment of purified SNS (Fig. 3B). These experiments confirm that the results obtained by direct sequence mapping of BrdU-labeled material represents a bona fide collection of active ORIs at heterochromatin and that TEs are a major source of ORIs in pericentromeric regions.

### Are TEs containing ORIs reactivated in cell cultures?

The activity of ORIs has been frequently associated with the expression level of the genomic loci where they are located (Sequeira-Mendes et al. 2009; Mechali et al. 2013). Although the expression of TEs is usually strongly repressed, some TEs can be activated under stress situations (Deragon et al. 2008; Lisch 2013). Notably, it was reported that in an Arabidopsis cell culture line typical heterochromatin marks change and some TEs are reactivated (Tanurdzic et al. 2008), in agreement with reports in Drosophila Kc and S2 cultured cells (Di Franco et al. 1992). Therefore, we determined the RNA levels across the ORI-containing region in each of the TEs selected previously. Our data showed that the RNAs derived from these elements were below detectable levels in all cases (Fig. S3). Similar results were obtained using either polyA-containing RNA or total RNA (Fig. S3). Furthermore, it is worth noting that the Athila elements, members of the Gypsy family, are among the most frequently reactivated TEs whereas the Atlantys elements, also from the Gypsy family, are very poorly reactivated (Tanurdzic et al. 2008). We found that ORIs colocalizing with Atlantys elements that account for ∼11% of all Gypsy elements are over-represented (43% of all ORIs in Gypsy elements). Consequently, we concluded that ORI-TE activity in our Arabidopsis cell culture line is independent of the transcriptional status of the TEs they are associated with. Based on these observations, we sought to identify whether a unique signature can be associated with the high preference of retrotransposon families for ORI specification.

### The activity of ORI-TEs is maintained with high levels of mC and is independent of G quadruplexes

The majority of ORIs colocalize with genes which, when highly expressed, tend to be highly methylated at CG positions within the gene body, but not at CHG or CHH, the other sequence contexts where C methylation is found in plants (Zhang et al. 2006). Moreover, the ±100 nt region around the ORI in euchromatin tends to be depleted of CG methylation (Costas et al. 2011), which suggests that ORI specification and activity may depend on low levels of methylation. TEs are heavily methylated in C residues of the three sequence contexts, and their methylation is actively maintained by RNA-directed DNA methylation (RdDM) and siRNAs (Matzke and Mosher 2014; Fultz et al. 2015). However, TEs may differ in their methylation state depending on the type, size or location (Ahmed et al. 2011; Zemach et al. 2013). Thus, we used the available methylation data of the Arabidopsis genome (Stroud et al. 2013) to ask whether differences in C methylation correlate with the preferential location of ORIs in certain TE family members. We found a tendency of Helitron elements, which do not colocalize with ORIs, to contain lower levels of C methylation for the three sequence contexts, whereas Gypsy elements, the most ORI-enriched TEs, showed higher methylation level (Fig. S4). This is in line with previous reports that showed that Helitrons tend to be less heavily methylated than Gypsy elements in Arabidopsis (Ahmed et al. 2011). Moreover, the level of C methylation of Gypsy elements does not vary depending on whether they colocalize or not with ORIs (not shown). Therefore, our data suggest that a low methylation level is not a requirement for ORI specification in TEs. Similar observations have been made for the heterochromatic X chromosome in mammalian cells where the level of C methylation does not affect ORI specification and usage (Gomez and Brockdorff 2004).

G quadruplexes (G4) have been frequently found in association with TEs (Kejnovsky et al. 2015) and with ORIs in mammalian cells (Besnard et al. 2012; Cayrou et al. 2012a; Valton et al. 2014; Comoglio et al. 2015). Thus, we also asked whether the presence of G4 was a determinant factor in the distribution of ORIs in Arabidopsis cells. We found first that G4 motifs are far more frequent in TEs than in genes whereas ORIs highly prefer a colocalization with genes. Second, most G4 motifs occur in a TE family known as ATREP18, which contains a canonical telomeric repeat (Cardenas et al. 2012) and that is also found in pericentromeric regions. This family is included within the annotation class “DNA/Other” that contains less than ∼1% of all ORI-TEs (Fig. S5). Third, and perhaps more relevant, both Gypsy and Helitron elements contain a very similar fraction of G4 motifs whereas they show an opposite preference to contain ORI-TEs (Fig. S5). Hence, our observations do not support the idea that G4 structures may be directly influencing ORI activity in Arabidopsis, and they do not explain the distribution of ORI-TEs among the different TE families found here.

### ORI-TE activity and the chromatin landscape

We next focused on the chromatin landscape around ORI-TEs to identify a possible common signature. We have previously shown that the entire Arabidopsis genome is characterized by nine different chromatin states (Sequeira-Mendes et al. 2014). To gain an overall view of the chromatin associated with ORI-containing TEs we looked for possible differences within TE families. We first investigated the whole chromatin signatures associated with the different TE families according to the known Arabidopsis chromatin states. Interestingly, we found that the majority of Gypsy, LINE and Copia families, which concentrate more than 80% of all ORI-TEs are associated with chromatin state 9 (Fig. 4A left pannels), which is characteristic of the GC-rich heterochromatin.

**Figure 4.**
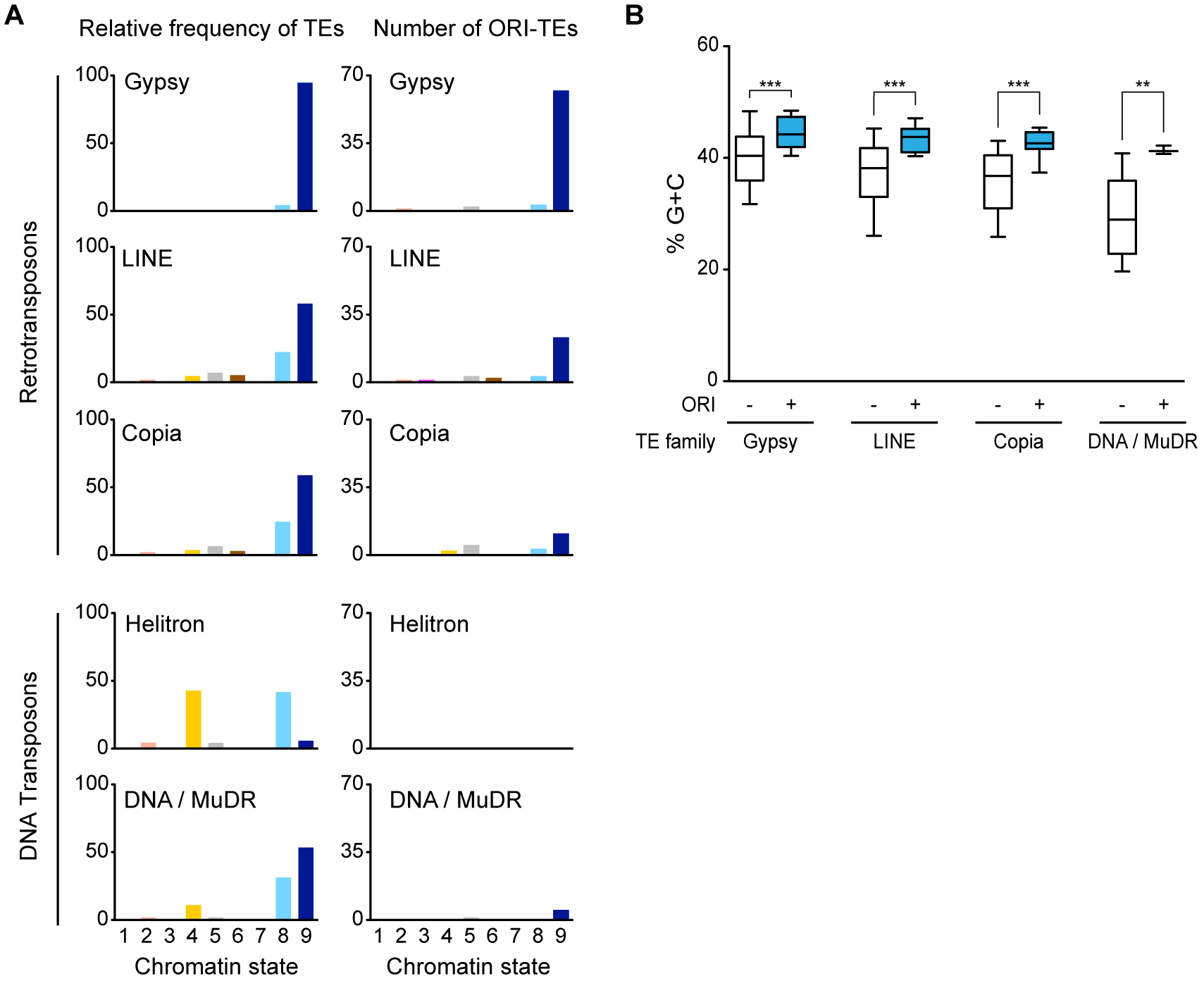
Distribution of retrotransposons and DNA transposons in the different chromatin states. (A) Relative frequency of several TE families (Gypsy, LINE, Copia, Helitron and DNA/MuDR), or ORI-TE of those families with respect to total nucleotide family content, in the nine chromatin states. Chromatin states, largely corresponding to various genomic elements, are as follows: state 1, TSS; state 2, proximal promoters; state 3, 5’ half of genes; state 4, distal promoters enriched in H3K27me3; state 5, Polycomb-regions; state 6, average gene bodies; state 7, long gene bodies; state 8, AT-rich heterochromatin; state 9, GC-rich heterochromatin. (B) Average G+C content of TEs with (blue) and without (white) ORIs in the different TE families. ***, p<0.0001; **, p<0.001 (unpaired t-test with Welch’s correction; wiskhers at 10-90 percentiles, outliers not represented in the graph).

This is particularly striking for the Gypsy elements, of which ∼95% are found in this heterochromatic state. On the contrary, Helitrons, which have a very low tendency to contain ORIs, are not associated with chromatin state 9 but to chromatin states 4 and 8 (Fig. 4A left panels). Chromatin state 4 is mainly associated with intergenic regions enriched in the Polycomb mark (H3K27me3), whereas chromatin state 8 is an heterochromatin state characterized by a lower GC content and a higher H3K27me3 level, as compared with the heterochromatin of chromatin state 9 (Sequeira-Mendes et al. 2014). Very interestingly, ORI-containing TEs tend to be in the chromatin state 9, independently of their family (Fig. 4A, right panel).

The main feature distinguishing the two heterochromatic states is the GC content, which is higher in chromatin state 9. In fact, this is a striking difference between TEs since the families that tend to contain ORIs (Gypsy, Copia and LINE) have a higher than genome average GC content. For instance, Gypsy elements contain 42.1% GC, the highest among TEs, compared with the 36.5% average GC content of the Arabidopsis genome (Fig. S6). On the contrary, Helitron elements are characterized by having a very low GC content (24.2%; Fig. S6). These differences in GC content do not have any impact on the potential to form G4 structures, as shown earlier, although they may have a direct impact on nucleosome organization (Liu et al. 2015; Zhang et al. 2015). Importantly, however, calculation of the average GC content of TEs that contain ORIs revealed that it was statistically significantly higher than in TEs of the same family that do not contain ORIs (Fig. 4B). This clearly suggests that a high GC content behaves as a determinant for ORI preference also at heterochromatic loci.

### ORI-TE activity is maintained with high H3K9me2 levels

The association of ORI-TEs with a heterochromatin state is somehow surprising as most ORIs are located within genes that colocalize with euchromatic marks found in very different chromatin states (Sequeira-Mendes et al. 2014). Even though we have already shown that transcription at ORI-TEs is not reactivated in cultured cells (Fig. S3) as chromatin may undergo changes in some cultured cells (Chupeau et al. 2013), we decided to analyze the chromatin marks associated with ORI-TEs in the Arabidopsis MM2d cultured cells.

We first looked at the overall levels of H3K9me2 and H3K27me1, two typical heterochromatic marks that strongly contribute to maintaining the silenced state of TEs in Arabidopsis (Law and Jacobsen 2010; West et al. 2014), by immunolocalization in cultured cells. H3K27me1 showed a pattern colocalizing with increased DAPI signal whereas H3K9me2 had a dotted appearance in nuclear sites enriched for H3K27me1 and DAPI positive regions (Fig. 5A), as it occurs in the nuclei of Arabidopsis plants. It must be noted that DAPI-stained chromocenters were not very apparent in nuclei of these cultured cells, suggesting a less condensed organization of the pericentromeric heterochromatin.

**Figure 5.**
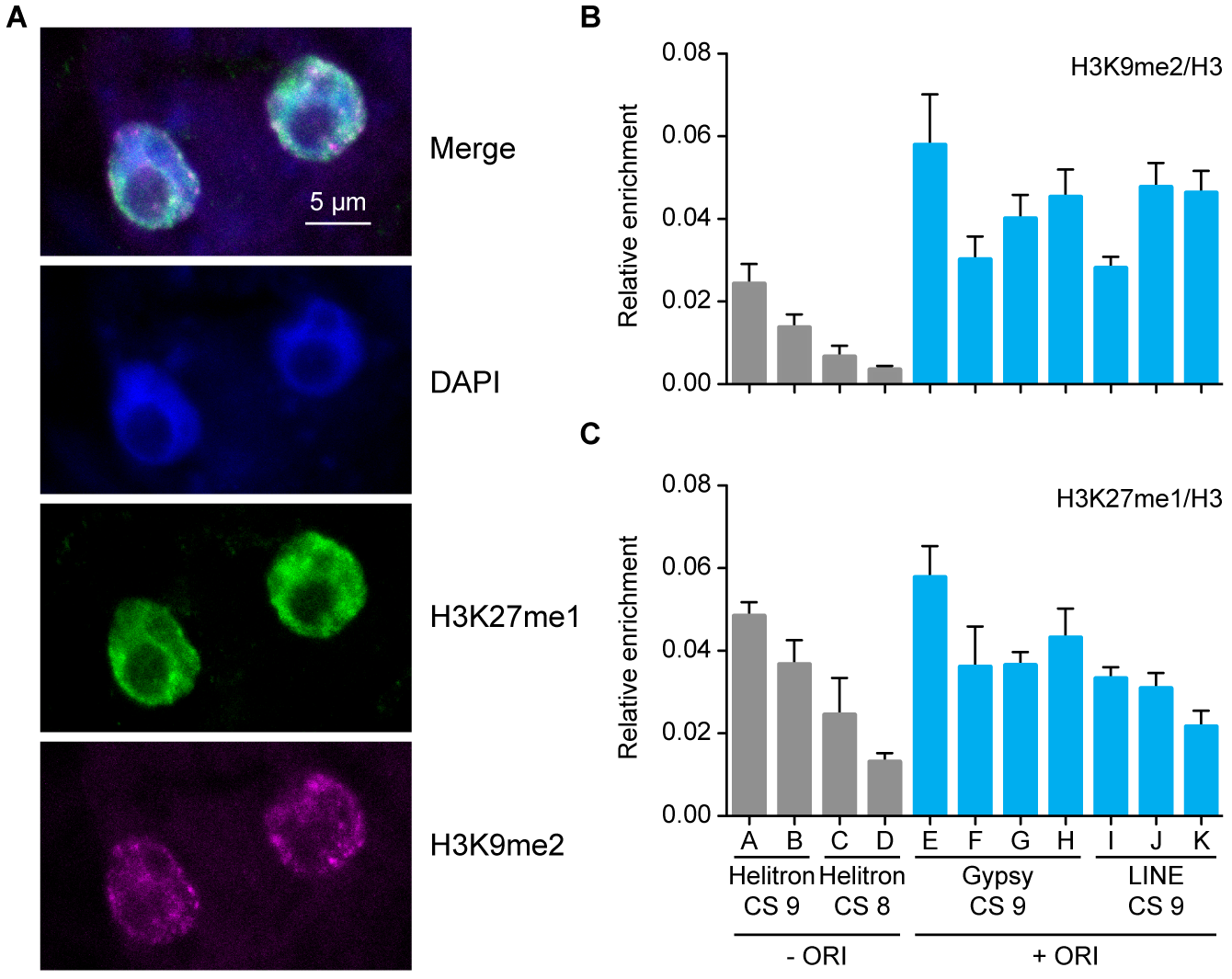
Heterochromatin marks in Arabidopsis MM2d cultured cells. *(A)* Immunolocalization of H3K9me2 (magenta) and H3K27me1 (green) in nuclei of cultured cells. Nuclei were stained with DAPI (blue). Levels of H3K9me2 *(B)* and H3K27me1 *(C)* determined by ChlP-qPCR in TEs representative of various families, chromatin states (CS) and with (blue bars) or without ORIs (grey bars). Enrichment values were made relative to the local H3 content determined by ChIP with anti-H3 antibody, as described in Methods. Two biological replicates and three technical replicates were evaluated. The mean values ± standard error of the mean is plotted. The codes for the primer pairs used to identify each TE, according to the list in Supplementary Table 5, are: A, AT2TE13970; B, AT4TE16735; C, AT2TE16335; D, AT4TE17050; E, AT4TE16726-2; F, AT4TE16726-3; G, AT1TE62820-3; H, AT1TE62820-5; I, AT2TE15565-2; J, AT2TE15565-3; K, AT4TE03295.

To determine more precisely the levels of H3K9me2 and H3K27me1 marks in cultured cells we performed ChIP and analyzed a subset of TEs containing a functional ORI. Although Helitron elements are not associated with ORIs, we also evaluated some Helitron elements located in the two heterochromatin states (AT-rich and GC-rich chromatin states 8 and 9, respectively). In all cases we normalized the measurements to the local H3 content determined by ChIP with anti-H3 antibody. We found that, in all the examples analyzed, the Gypsy and LINE elements (GC-rich heterochromatin state 9) contain a high level of H3K9me2 (Fig. 5B). We also found that in general the H3K9me2 level was higher in retrotransposons than in Helitron elements, independently of their chromatin state (Fig. 5B), similar to what was reported in maize (West et al. 2014). In the case of H3K27me1, which is typical of heterochromatin and crucial to prevent re-replication (Jacob et al. 2010), ChIP experiments revealed that the TEs analyzed showed various levels of H3K27me1 independently of (i) being Gypsy, LINE or Helitron, (ii) their chromatin signature and (iii) their colocalization with ORIs (Fig. 5C). Alterations in the nuclear DNA content are indicative of massive defects in re-replication control and, indirectly, of possible decrease in H3K27me1, as it occurs in the *atxr5,atxr6* mutant (Jacob et al. 2010). Consistent with our ChIP data, we could not detect any significant alteration in the DNA content profile of cultured Arabidopsis cells (Fig. S7). Since retrotransposons are enriched for ORIs and H3K9me2 and there is a lack of correlation of H3K27me1 with ORI-TEs, these marks seem to be unrelated to ORI activity.

## Discussion

The results presented here show that whereas in euchromatic regions ORIs are almost exclusively located within genes, in the heterochromatic pericentromeric regions a significant fraction of ORIs colocalizes with TEs. This underscores the relevance of retrotransposons in contributing to genome replication, a key process during the cell cycle. We show here that the epigenetic marks associated with ORI-TEs (high methylation at all cytosine contexts, H3K9me2 and H3K27me1) are typical of heterochromatin and very different from those associated with euchromatic ORIs, suggesting that these marks do not interfere with ORI specification. Interestingly, ORI-TEs are not randomly distributed among TEs and show a striking tendency to colocalize with retrotransposons, and in particular with Gypsy elements. Transcription is the first and obligate step for mobilization of all retrotransposons, whereas DNA transposons are mobilized by a DNA intermediate and don’t need to be transcribed. This makes retrotransposons more similar to genes than any other TE. Indeed, whereas most retrotransposons are silent in most plant tissues, their activation under stress or in particular mutant backgrounds confirms that they retain the capacity to be transcribed and to transpose (Bucher et al. 2012; Cavrak et al. 2014). However, activity of ORI-TEs cannot be explained by transcription through TE sequences. In addition, both genes and retrotransposons show an above average GC content, which makes their sequences different from most DNA transposons and particularly Helitron elements. Importantly, TEs with ORIs possess a higher GC content than TEs without ORIs, independently of their TE family. Therefore, these results lead us to propose that a high local GC content, typical of the heterochromatin state 9 where the vast majority of ORI-TEs are located, in combination with the potential to be transcribed, characteristic of the genomic organization of retrotransposons, are the major features of ORIs colocalizing with TEs. These characteristics allow certain TE families to contribute to a significant fraction of ORIs in heterochromatic regions. This can be crucial to ensure correct replication of heterochromatic domains, which have a low gene density, thus compensating for the high preference of ORIs to localize in genes.

## Methods

### Plant material and growth conditions

*Arabidopsis thaliana* MM2d cell line (Menges and Murray 2002) was grown at 26 °C and 120 rpm, in the absence of light. The cells were subcultured every 7 days into fresh Murashige & Skoog medium (MS, pH 5.8, Duchefa) supplemented with 3% sucrose (Duchefa), 0.5 ug/mL 1-naphthaleneacetic acid (Duchefa), 0.1 ug/mL kinetin (Sigma) and 0.103 ug/mL vitamins (Duchefa).

### BrdU sequencing data analysis

BrdU sequencing data reads (GEO GSE2182; (Costas et al. 2011)) were trimmed down to 50 nt from the 3’ end and mapped to the reference Arabidopsis genome (TAIR10) using BOWTIE aligner (Langmead et al. 2009), allowing up to three mismatches and discarding multihit reads. PCR duplicate reads were removed using an in-house script. Peak calling was performed using MACS1.4 (Zhang et al. 2008) with a cutoff value of 10^-6^. Neighboring peaks were merged when interpeak distance was less than 260 nt. Peaks smaller than 200 nt were removed from the analysis. Analogous results were obtained when using a similar (Spyrou et al. 2009) algorithm (data not shown). The same analysis was carried out using only the multihit reads.

### ORI distribution and classification

General annotation coverage was calculated with the complete set of annotations from TAIR10, discarding "transposon_fragment" as it is redundant with the "transposable_element" annotation. Pericentromeric regions were defined as the regions where the gene coverage in 1 Mb bin was equal or lower to 40%. ORIs were attributed to a type of annotation (genes, TEs or particular TE families) only for unambiguous non-overlapping annotation. TE family coverage was calculated within the TE genome space (total TE nucleotide content).

### C methylation, G quadruplex, GC content and chromatin states analysis

CG, CHG and CHH methylation data were retrieved from (GEO GSE39901) (Stroud et al. 2013). The presence of G quadruplexes in the Arabidopsis genome was predicted using the Quadparser software (Hershman et al. 2008) allowing a spacing of 7 nt between G strings. The GC content of the genome was calculated in bins of 50 nt. For the analysis of the distribution of TE among the different chromatin states (Sequeira-Mendes et al. 2014), the relative frequency of each TE family in each state was determined by the coverage of the family in that particular state relative to the total coverage of the TE family in the genome. For the distributions of ORI-TEs among the different chromatin states the ORI midpoint was considered. All the bioinformatics analyses were performed with in-house Perl scripts and BEDtools suite utilities (makewindows, genomecov, merge, intersectBed) (Quinlan 2014).

### Cell synchronization

Cells on exponential phase (4 days after subculture) were synchronized in G0/G1 by growing them in MS without sucrose for 24 hours. To release the cell cycle block the medium was replaced with MS with sucrose (Menges and Murray 2002). Samples for analysis were taken at 2 (G1/S transition), 3.5 (early S) and 7 (late S) hours.

### Isolation of Short DNA Nascent Strands (SNS)

The short replication intermediates used in the ORI activity qPCR assays were purified essentially as described (Costas et al. 2011). At day 4 after passage, 100 mL of the asynchronous cell suspension were either directly collected for SNS preparation or synchronized at the desired time points (2, 3.5 or 7 h) before SNS isolation.

### RNA analysis

Total RNA from asynchronous cells was isolated at day 4 after subculture using Trizol reagent (Invitrogen) according to manufacturer’s instructions. Total RNA was treated with DNase I (Roche) and 1 ug was reverse-transcribed with Superscript III (Invitrogen) using an oligo-dT primer (mRNA) or random hexamers (total RNA). Two microliters of a 3-fold diluted cDNA reaction were used as template in qPCR, and the primers listed in Table S5.

### Immunolocalization

MM2d cells were collected at 4 days after subculture and fixed in 4% paraformaldehyde in microtubules stabilizing buffer (MTSB; 50 mM PIPES, pH 6.9, 5 mM EGTA, 5 mM MgSO_4_), for 10 min plus 5 min with vacuum infiltration. Cells were washed with MTSB, PBS and water and air-dried on superfrost plus slides (Thermo Scientific). Cells were re-fixed in 4% paraformaldehyde in MTSB for 30 min and washed with MTSB. Cell wall was partially digested with 20 mg/mL driselase (Sigma) in MTSB for 45 min at 37 °C and the slides were washed with PBS. Membranes were permeabilized with 10% DMSO, 3% Igepal CA-630 in MTSB for 1 h. Non-specific sites were blocked in 3% BSA, 10% Horse Serum (HS) in PBS for 1 h at 37 °C. H3K9me2 and H3K27me1 were detected with antibodies (Abeam ab1220 and Millipore 07-448, respectively) diluted 1:1000 in 1% BSA, 10% HS, 0.1% Tween-20 in PBS at 4 °C overnight. Slides were washed with 3% BSA in PBS and incubated with donkey anti-mouse 555 and anti-rabbit 488 (A-31570 and A-21206 Thermo Scientific, respectively) diluted 1:500 in 1% BSA, 10% HS, 0.1% Tween-50 in PBS for 1 h. Following washes in 3% BSA in PBS, nuclei were counterstained with DAPI (Merck), washed with PBS and mounted in Mowiol 4-88 (Sigma). The localization of H3K9me2 and H3K27me1 in immunostained cells was analyzed by confocal microscopy (LSM710 Zeiss). Images were processed using Fiji.

### Chromatin immunoprecipitation

MM2d cells were harvested 4 days after subculture and fixed using ice-cold 1% formaldehyde in PBS and applying vacuum infiltration (3 rounds of 6 min on/4 min off). The cross-linking was stopped by the addition of 0.125 M glycine, infiltrating for another 5 minutes. The grinded material was resuspended in Extraction Buffer (0.25 M sucrose, 10 mM Tris-HCI, pH 8.0, 10 mM MgCI2, 1% Triton X-100, 1 mM PMSF, and protease inhibitor cocktail for plant cell extracts (Sigma)). Nuclei were pelleted by centrifugation, resuspended in Lysis Buffer (50 mM Tris-HCI, pH 8.0, 10 mM EDTA, 1% SDS, 1 mM PMSF, and protease inhibitor cocktail) and disrupted by sonication in a Bioruptor Plus (Diagenode) for 30-45 cycles of 30 s on and 30 s off, at high power mode. One microgram of soluble chromatin was employed per ChIP reaction, using the following antibodies: anti-H3K9me2 (Abcam ab1220, 3 ug), anti-H3K27me1 (Millipore 07-448, 1 ug), anti-total H3 (Abcam ab1791, 2 ug), or anti-rat IgG (Abcam ab6703, 2 |jg) as a negative control. Immune complexes were recovered with 50 uL of protein G agarose beads (SCBT) and washed and eluted essentially as described (Villar and Kohler 2010).

All qPCRs (SNS, cDNA, and ChIP) were performed using GoTaq Master Mix (Promega) according to the manufacturer’s instructions in an ABI Prism 7900HT apparatus (Applied Biosystems) using the primers listed in Table S5.

### Flow cytometry

MM2d cells were collected at either 4 or 7 days after subculture by vacuum filtration and the retentate was chopped in Galbraith solution (45 mM MgCI_2_, 20 mM MOPS, 30 mM sodium citrate, 0.1% Triton X-100, pH 7.0). Nuclei were filtered through a 30-um nylon net filter (Millipore) and stained with 2 ug/mL DAPI. Nuclei populations were analyzed using a FACSCanto II High Throughput Sampler cytometer (Becton Dickinson) and FlowJo v10.1rS software (FlowJo).

## Supplemental files

Figures S1 to S7.

Tables S1 to S5.

## Acknowledgments

We thank the Confocal Microscopy and Flow Cytometry Services of CBMSO and V. Mora-Gil for technical assistance. We also thank Martí Bernadó of CRAG’s genomics facility for assistance with statistical analyses. We thank E. Martinez-Salas for comments on the manuscript. Z.V. has been recipient of a FPI Predoctoral Fellowship from MINECO. This research was supported by grants BFU2012-34821 (MINECO), BIO2013-50098-EXP (MINECO) and BFU2015-68396-R (MINECO/FEDER) to C.G., and grants BFU2009-11932 and AGL2013-43244-R to J.M.C. from MINECO, and by an institutional grant from Fundación Ramón Areces to the Centro de Biologia Molecular Severo Ochoa.

### Author contributions

C.G. and J.M.C. conceived the study. Z.V. together with J.S.-M. carried out functional analysis of ORIs, the ChlPs, expression and cellular analysis. C.C. participated in the early stages of ORI analysis. R.P. reanalyzed the BrdU sequencing data. J.M., with the initial participation of E.H., did the bioinformatics analysis of ORI-TEs, TE families and chromatin features. All authors discussed the results. C.G. and J.M.C. wrote the manuscript with the participation of all authors, who agree on the final text.

### Competing interests

Authors declare that they have no competing interests

